# A Scalable Screening of *E. coli* Strains for Recombinant Protein Expression

**DOI:** 10.1101/2021.03.09.434560

**Authors:** Luana G. Morão, Lívia R. Manzine, Lívia Oliveira D. Clementino, Carsten Wrenger, Alessandro S. Nascimento

## Abstract

Structural biology projects are highly dependent on the large-scale expression of soluble protein and, for this purpose, heterologous expression using bacteria or yeast as host systems is usually employed. In this scenario, some of the parameters to be optimized include (*i*) those related to the protein construct, such as the use of a fusion protein, the choice of an N-terminus fusion/tag or a C-terminus fusion/tag; (*ii*) those related to the expression stage, such as the concentration and selection of inducer agent and temperature expression and (*iii*) the choice of the host system, which includes the selection of a prokaryotic or eukaryotic cell and the adoption of a strain. The optimization of some of the parameters related to protein expression, stage (*ii*), is straightforward. On the other hand, the determination of the most suitable parameters related to protein construction requires a new cycle of gene cloning, while the optimization of the host cell is less straightforward. Here, we evaluated a scalable approach for the screening of host cells for protein expression in a structural biology pipeline. We evaluated four *Escherichia coli* strains looking for the best yield of soluble heterologous protein expression using the same strategy for protein construction and gene cloning and comparing it to our standard strain, Rosetta 2 (DE3). Using a liquid handling device (robot), *E. coli* pT-GroE, Lemo21(DE3), Arctic Express (DE3), and Rosetta Gami 2 (DE3) strains were screened for the maximal yield of soluble heterologous protein recovery. For the genes used in this experiment, the Arctic Express (DE3) strain resulted in better yields of soluble heterologous proteins. We propose that screening of host cell/strain is feasible, even for smaller laboratories and the experiment as proposed can easily be scalable to a high-throughput approach.

## 1. Introduction

Many drug discovery projects rely on the structure determination of the drug targets. Indeed, to the best of our knowledge, the so-called ‘structure-based rational design’ of drug candidates dates to 1976, when Beddell and coworkers designed a series of compounds to fit the human haemoglobin site [1]. Typical structural biology pipelines require pure and soluble protein in high amounts, which is usually achieved by heterologous expression in *Escherichia coli* [2,3] or in yeast cells, such as *Pichia pastoris* or *Saccharomyces cerevisae*, or even in fungus, such as *Aspergillus niger* (e.g., [4]).

*E. coli* is the host cell of choice in most structural biology approaches, due to the ease in handling, rapid doubling time, and very well-established protocols for expression. In our experience, protein expression in *E. coli* for high-throughput (HT) pipelines resulted in yields of up to 946 mg of protein expressed per liter of culture, with an average of about 60-70 mg.L^-1^ for enzymes of the glycoside hydrolase (GH) superfamily [5]. Although this pipeline works very well for most cases, some ‘*difficult cases*’ require additional efforts, including new cloning strategies, such as the addition of fusion tags or even change of tags from C-terminus to N-terminus and vice versa [6]; the exchange of expression vector, suiting of the expression protocol or replacing the host cell.

In this context, several HT strategies have been developed for parallel cloning of genes of interest in different constructions, which explore different fusion proteins, different tags, and the location of the tag [6–8]. Also, protocols for increasing the yield of soluble expression of target proteins are now abundant in the literature (e.g., [3]). Finally, a myriad of commercial *E. coli* strains is available for efficient heterologous protein expression, promising high yield and enhanced solubility, especially for insoluble protein, making the choice for a particular strain a non-straightforward task.

Here we show the results of a scalable parallel screen for the expression of difficult cases using four *E. coli* strains, and the same strategy for protein construction and gene cloning. The aim of this study was to develop a comparison of the expression levels of proteins of interest (real-scenario cases) in different *E. coli* strains in a HT format. The assay can be readily scalable for an HT format and also include more strains. We show that the screening for expression efficiency in different strains can be a feasible and rapid way for the ‘salvage’ of target proteins that failed to be soluble expressed in the first HT approach, avoiding going back to the cloning stage of the structural biology pipeline. An automated liquid handling device was central to the screening, allowing rapid analysis of the expression in 96-well plates.

## 2. Materials and methods

All the genes tested in this work were cloned in the pET-Trx1A/LIC using the strategy previously described by Camilo & Polikarpov [5]. Besides, for a particular target (target **T16**) the pET-NESG vector was used. The pET-Trx1A/LIC is a ligation independent cloning-based vector where the target gene is expressed with an N-terminal His-tag fused to thioredoxin and a cleavage site for the Tobacco Etch Virus (TEV) protease.

### 2.1. Gene Cloning

The LIC cloning protocol using pET-Trx1A/LIC vector was performed as described previously [5]. Briefly, the plasmid was linearized by PCR using the following oligonucleotides: Fw-5’ TGGCGCCCTGAAAATAAAG; Rv-5’CCGCGTCGGGTCAC and, for targets, genomic DNA were used as templates in PCR amplification using Phusion High-fidelity DNA Polymerase (New England Biolabs). PCR products were individually treated for 16 h at 37 °C with 20 units of DpnI enzyme (New England Biolabs). Following gel extraction using Wizard SV Gel and PCR CleanUp System (Promega), 500 ng of purified and linearized vector and 200 ng of each specific gene were treated with T4 DNA polymerase enzyme (Fermentas) in the presence of 2.5 mM dTTP (vector) and 2.5 mM dATP (gene) for 30 min at 22 °C. Annealing of LIC vectors and insert was performed after incubation of 3 µL of T4 Polymerase-treated PCR fragments with 1 µL of T4 Polymerase-treated vector at 25 °C for 30 min. Then, the mixture was used for *E. coli* DH5αcompetent cells transformation and positive clone selection achieved in culture plates containing LB agar medium with 50 µg mL^-1^ kanamycin after overnight incubation at 37 °C.

### 2.2. Protein Expression and Analysis

Sixteen target genes were cloned following the procedures above. The resulting vectors were used for *E. coli* expression in four DE3 strains: Lemo21, pT-GroE, Rosetta-Gami 2, and Arctic Express. Transformations were performed in 24-well-LB-agar plates containing: kanamycin (50 µg mL^-1^) and chloramphenicol (34 ug mL^-1^) for Lemo21, pT-GroE, and Rosetta-Gami 2 cells; kanamycin (50 µg mL^-1^) and gentamycin (20 µg mL^-1^) for Arctic Express. Colonies were inoculated in DW96 preculture plates containing 1 mL of LB medium supplied with cells specific antibiotics, sealed, and incubated overnight at 37 °C with shaking. Then, 100 µL of preculture was inoculated in 5 DW24 plates containing 4 mL of LB with specific antibiotics (plus 0.5 mM rhamnose for Lemo21 cells) and grown at 37 °C under 150 rpm agitation for 5 h. Afterward, aliquots were collected and IPTG induction was performed using 0.4 mM final concentration for Lemo21 and 1 mM for the other strains. The temperature was reduced to 18 °C (pT-GroE and Rosetta-Gami 2), 30 °C (Lemo21), or 11 °C (Arctic Express), and expression was conducted for 16 h at 150 rpm. In parallel, 100 µL of pre-culture was inoculated in 5 DW24 plates containing 4 mL of ZYP5052 [9] auto-induction medium, and the expression was performed as LB-conditions without IPTG. The plates were centrifuged for 10 min at 2000*x*g and each pellet was resuspended with 1 mL of lysis buffer (Tris–HCl 50 mM, NaCl 300 mM, lysozyme 0.25 mg mL^-1^, 0.1 mM PMSF, at pH 8). Cells were lysed by freezing at -80 °C overnight and thawing at 17 °C for 1 h under agitation. DNAseI (Sigma) was added at 10 µg mL^-1^ final concentration following incubation at 17 °C for 15 min. After centrifugation at 3700 rpm, the lysate purification was carried out in an automated platform Freedom EVO 200 (Tecan) using Ni-Sepharose 6 Fast Flow Resin (GE) in 96-well Receiver Plates 20 µm (Macherey–Nagel). The lysate passed through the resin by vacuum filtration followed by resin washing with 1 mL of buffer A (Tris–HCl 50 mM, NaCl 300 mM, pH 8) and buffer B (Tris-HCl 50 mM, NaCl 300 mM, imidazole 20 mM, pH 8) for removal of unspecific and unbounded proteins. Elution of interest proteins was performed with 150 µL of buffer C (Tris-HCl 50 mM, NaCl 300 mM, imidazole 300 mM, pH 8) and samples were analyzed by gel electrophoresis for expression protein screening evaluation. The expression yield of the soluble proteins expressed was assessed by Coomassie-stained SDS-PAGE 15%, after purification by nickel affinity chromatography (HTS). Thus, the bands observed in the gels are due to soluble proteins that were purified by nickel affinity chromatography. An overall view of the entire pipeline, from cloning to the analysis of heterologously expressed protein is shown in Figure 1.

**Figure 1.**
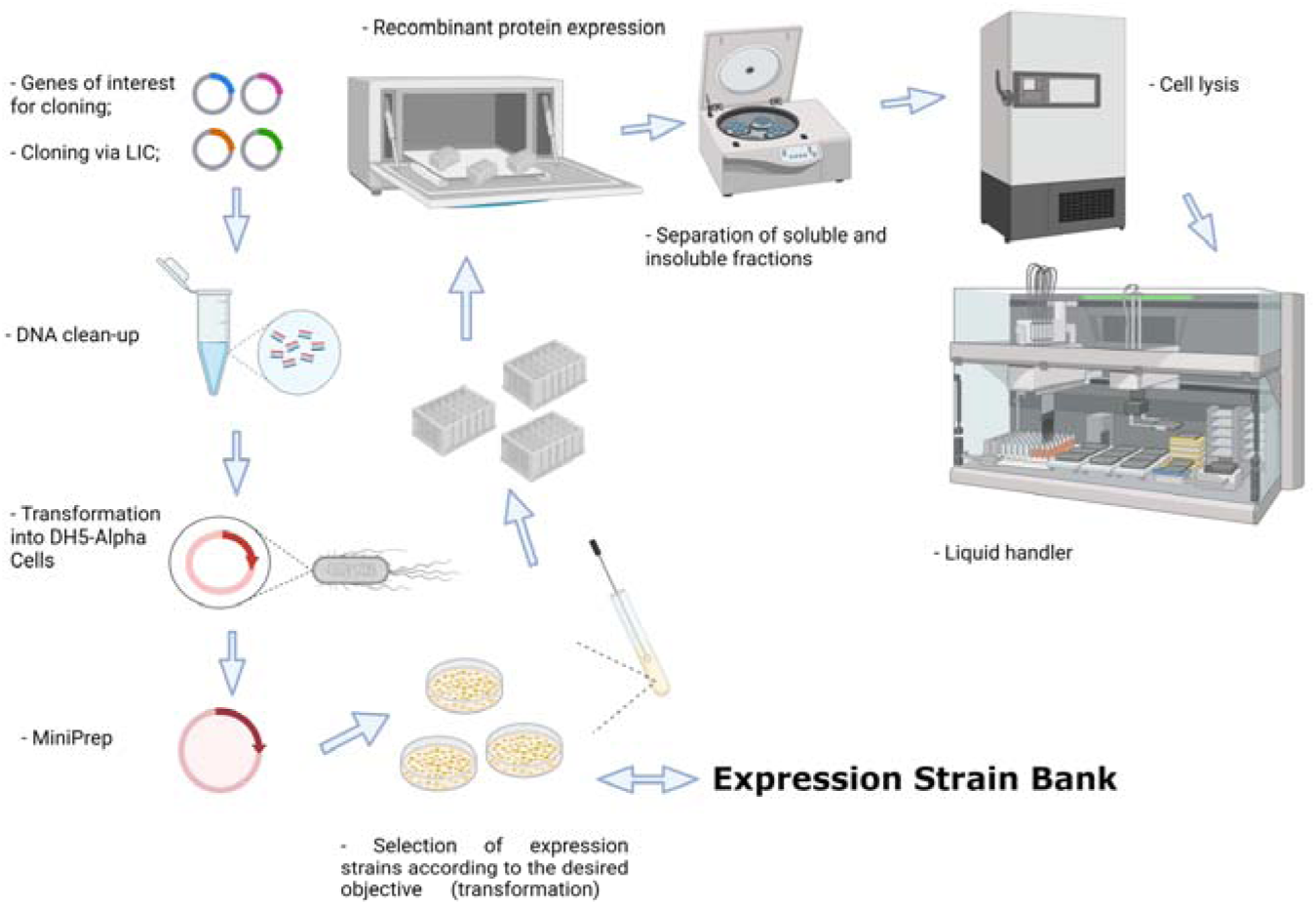
Overview of the pipeline used in this work. Selected genes are cloned in the pET-Trx1A/LIC vector. After plasmid propagation in *E. coli* DH5α cells, the purified vector is used for transformation of a set of *E. coli* strains, the Expression Strain Bank. The cells are cultured in 96-well plates in auto-induction medium and, after cell lysis by freezing, a liquid handling robot separates the soluble fraction, incubates with the affinity resin, washes, and elutes the purified targets, which are further analyzed by SDS-PAGE 15%. Created with BioRender.com.

The intensities of the SDS gel bands were quantified using the software ImageJ [10]. Since the intensities of the gels might change from one gel to another due to changes in staining or destaining, for example, the raw intensities were normalized for the intensity of the molecular marker at 30 kDa in each gel. Since the markers are used in the same amount and from the same vendor, it provides an absolute scale for internal normalization. The band intensities are reported in fraction of the intensity of the reference band (molecular marker ar 30 kDa) and in a semi-quantitative scale, from ‘+’ to ‘++++’, according to the normalized intensities computed with ImageJ.

For the targets **T11** and **T14**, two glycosyltransferases, different parallel strategies were used. Besides the protocol previously described, three alternative approaches were attempted for **T11**: two different truncated constructs, removing different portions of the C-terminus of the protein (**T12** and **T13**), and the expression using the vector pET-NESG (**T16**). For T14 a C-terminus truncated form was also evaluated (**T15**).

## 3. Results

Sixteen ‘difficult cases’ genes were identified in our typical protein expression pipeline. In this pipeline, the genes are cloned in the pET-Trx1A/LIC vector and expressed in *E. coli* Rosetta 2 (DE3) strain. In 13 out of the 16 cases, no soluble protein expression could be observed, or the expression yield was very low for a structural biology pipeline. The exceptions were targets **T3, T5** and **T7**. The targets include bacterial (*Staphylococcus aureus, Enterococcus faecalis*, and *Mycobacterium tuberculosis*) enzymes, as well as plasmodial enzymes (*Plasmodium falciparum* and *Plasmodium vivax*). The genes were initially synthesized with codon optimization for *E. coli* expression. The only exception was the *E. faecalis* genes. These genes were cloned directly from the genomic DNA (DMS20478). All the protein products present a molecular weight of 30 to 50 kDa and some of them are under-represented in the PDB, suggesting that they are difficult cases for protein expression in the structural biology pipeline in general. A summary of the genes selected for the screening of soluble expression in *E. coli* strains is shown in Table 1.

**Table 1.** Target genes used for protein expression. The isoelectric point and molecular mass were computed with the ProtParam server [14]. TBP = thiamine biosynthesis pathway. PBP = pyridoxal 5’-phosphate biosynthesis pathway. GT= glycosyltransferase. Targets flagged with * are where some soluble expression was also observed in *E. coli* Rosetta 2.

| Target | Pathway | Organism | N<sub>res</sub><sup>a</sup> | N<sub>Cys</sub><sup>a</sup> | pI<sup>a</sup> | MW<sup>b</sup> (kDa) |
| --- | --- | --- | --- | --- | --- | --- |
| **T1** | TBP | *E. faecalis* | 276 | 1 | 5.47 | 43.3 |
| **T2** | TBP | *S. aureus* | 276 | 2 | 5.78 | 44.2 |
| **T3\*** | PBP | *M. tuberculosis* | 299 | 1 | 5.24 | 45.4 |
| **T4** | PBP | *M. tuberculosis* | 199 | 2 | 5.33 | 35.2 |
| **T5\*** | PBP | *P. vivax* | 302 | 7 | 6.62 | 47.2 |
| **T6** | PBP | *P. vivax* | 219 | 7 | 6.54 | 38.5 |
| **T7\*** | TBP | *P. falciparum* | 310 | 16 | 6.18 | 49.0 |
| **T8** | TBP | *P. falciparum* | 302 | 11 | 8.88 | 47.9 |
| **T9** | TBP | *P. falciparum* | 400 | 13 | 8.89 | 60.7 |
| **T10** | PBP | *P. falciparum* | 219 | 9 | 6.43 | 38.6 |
| **T11** | GT | *E. faecalis* | 241 | 2 | 6.23 | 40.9 |
| **T12** | GT | *E. faecalis* | 210 | 2 | 6.22 | 37.7 |
| **T13** | GT | *E. faecalis* | 130 | 1 | 4.39 | 28.4 |
| **T14** | GT | *E. faecalis* | 325 | 0 | 9.08 | 51.2 |
| **T15** | GT | *E. faecalis* | 220 | 0 | 5.59 | 39.1 |
| **T16** | GT | *E. faecalis* | 241 | 2 | 6.23 | 26.9 |

<sup>a</sup> Applies to the target sequence, without considering the fusion protein sequence.
<sup>b</sup> Molecular mass computed with the fusion tag.

After purification, the obtained expression levels were compared by SDS-PAGE and image analysis with ImageJ. Figure 2 shows the electrophoresis analysis obtained for the expression of the target genes and the results are also summarized in Table 2. Targets **T1, T2**, and **T7** are orthologous enzymes from *E. faecalis, S. aureus*, and *P. falciparum*, respectively, and all of them showed negligible soluble expression in *E. coli* Rosetta 2 strain. When tested for the expression in Rosetta Gami 2, pT-GroE, Lemo21, and Arctic Express, we observed soluble recovery of **T1** in Lemo21, Rosetta Gami 2, Arctic Express, and, with a smaller yield, pT-GroE. Additionally, **T2** was recovered in minimal amounts in Arctic Express, while **T7** was recovered in higher amounts in the same strain.

**Table 2.** Summary of the soluble protein recovery using different *E. coli* strains culture medium. (left) The expression levels were quantified using the software ImageJ and normalized by the intensity of the band from the molecular marker at 30kDa, as shown in Figure 2. (right) The normalized intensities shown in the left are reported in the scale no expression (-, intensity I < 10%), small yield (+, 10% < I < 20%), reasonable yield (++, 0.2 < I < 0.4), good yield (+++, 0.4 < I < 0.6) and excellent yield (I > 0.6).

| Target | Arctic Express (DE3) | Rosetta Gami 2 (DE3) | pT_GroE (DE3) | Lemo 21 (DE3) | Target | Arctic Express (DE3) | Rosetta Gami 2 (DE3) | pTGroE (DE3) | Lemo 21 (DE3) |
| --- | --- | --- | --- | --- | --- | --- | --- | --- | --- |
| T1 | 28% | 32% | 11% | 48% | T1 | ++ | ++ | + | +++ |
| T2 | 15% | 4% | 1% | 1% | T2 | + | - | - | - |
| T3 | 58% | 6% | 23% | 8% | T3 | +++ | - | ++ | - |
| T4 | 1% | 1% | 0% | 0% | T4 | - | - | - | - |
| T5 | 22% | 8% | 1% | 16% | T5 | ++ | - | - | + |
| T6 | 135% | 49% | 0% | 28% | T6 | ++++ | +++ | - | ++ |
| T7 | 28% | 4% | 6% | 8% | T7 | ++ | - | - | - |
| T8 | 22% | 0% | 1% | 1% | T8 | ++ | - | - | - |
| T9 | 2% | 28% | 13% | 1% | T9 | - | ++ | + | - |
| T10 | 38% | 62% | 37% | 75% | T10 | ++ | ++++ | ++ | ++++ |
| T11 | 1% | 1% | 4% | 1% | T11 | - | - | - | - |
| T12 | 18% | 5% | 4% | 1% | T12 | + | - | - | - |
| T13 | 57% | 5% | 17% | 1% | T13 | +++ | - | + | - |
| T14 | 11% | 1% | 6% | 1% | T14 | + | - | - | - |
| T15 | 59% | 1% | 12% | 1% | T15 | +++ | - | + | - |
| T16 | 1% | 1% | 11% | 4% | T16 | - | - | + | - |

**Figure 2.**
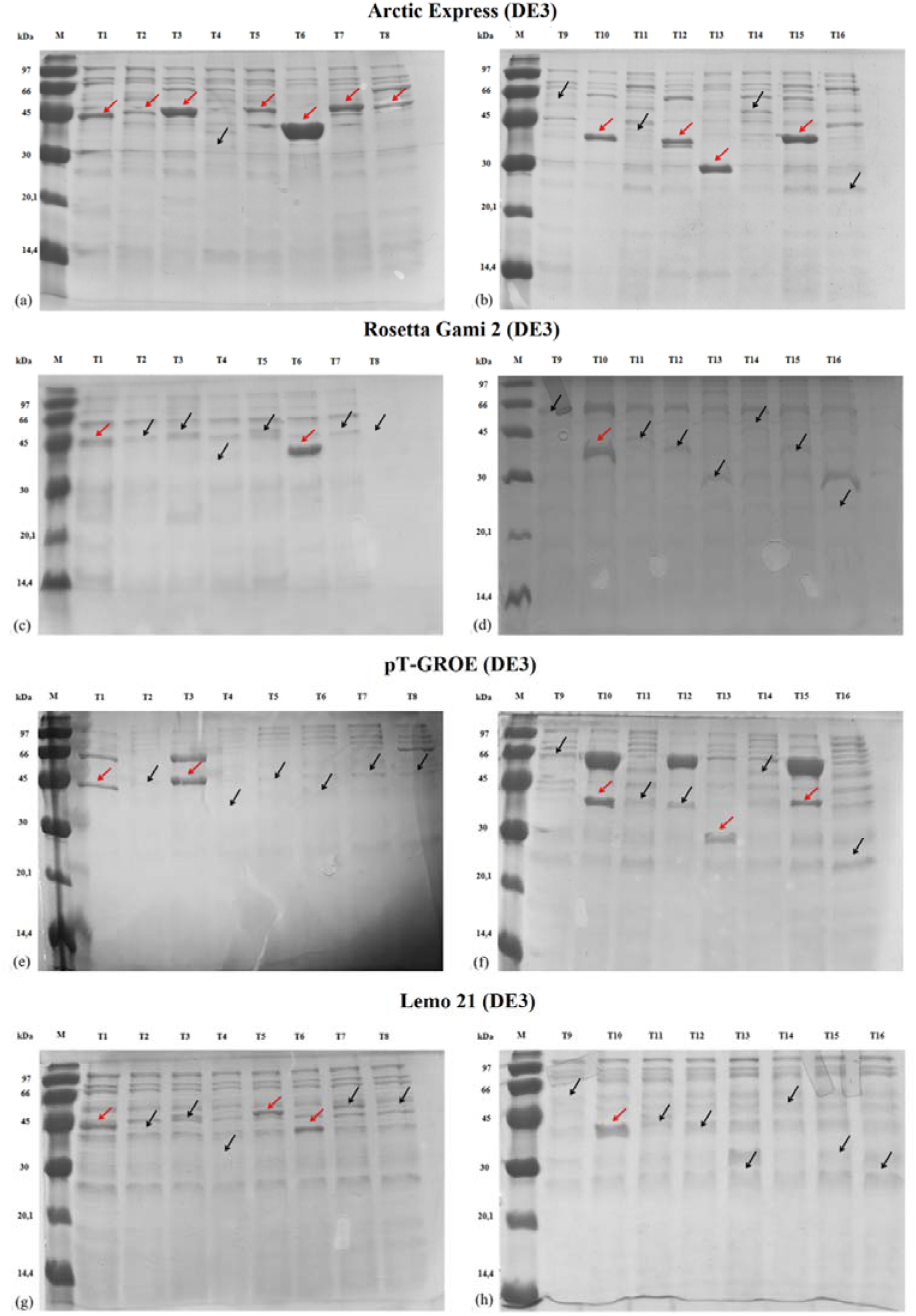
SDS-PAGE (15%) analysis of soluble protein recovery after affinity chromatography in the four strains used in this work. The red arrows indicate the recovered protein with the correct molecular weight. The black arrows indicate the molecular weight of the proteins that were not recovered or only minimally recovered.

The targets **T3, T4, T5, T6**, and **T10** are five enzymes of the same pathway in *M. tuberculosis* and *Plasmodium* species. Out of these five targets, the Arctic strain recovered four of them in the soluble form, while pT-GroE recovered two of them with a good expression pattern (**T3** and **T10**). Lemo21 strain recovered **T5, T6**, and **T10**, and Rosetta Gami 2 recovered targets **T6** and **T10**. The target **T9** was recovered in very low yields in the pT-GroE strain and to a greater extent in Rosetta Gami 2, while the Arctic Express strain recovered reasonable amounts of the target **T8**.

The targets **T11** and **T14** are glycosyltransferases, which are targets known to be insoluble in many cases. **T11** was also tested for its expression in two truncated forms (**T12** and **T13**) and also in the full-length protein in the pET-NESG vector (**T16**). A truncated form of **T14** was also tested in the pET-Trx1A/LIC vector and is listed as **T15**. The strains Lemo21 and Rosetta Gami 2 were not able to recover any of the targets. The pT-GroE strain recovered three of the truncated forms for these targets (**T13, T15, and T16**) in very small amounts. Finally, the Arctic strain recovered **T13** and **T15** in reasonable amounts and targets **T12** and **T14** in smaller amounts, with very good contrast to the background proteins observed in the SDS-PAGE.

## 4. Discussion

The protein purification procedures, as described in the Materials and Methods section, are assisted by a liquid-handling device. This automation device is central to the protocol used in this work, allowing a rapid evaluation of the amount of soluble protein obtained in cell lysis. Also, the automated procedure is readily scalable for an increased number of targets or an increased number of strains tested. The procedure followed the same HT strategy previously established [5]. The actual bottleneck was the protein expression since each *E. coli* strain has its typical resistance marker and its optimal temperature for expression. In our assay, the growing temperature was set to 37 °C for all strains and different temperatures were used in the induction stage, by allocating each strain to a different shaker. Although this step requires increased human effort, it is easily scalable to include more strains, in particular, if the additional strains can be used in the same expression temperatures, or even for more genes, since they can be used in the same plate in each shaker. After the protein expression step, the samples were collected on a single plate for the purification procedure. The process can be further optimized for an HT format, but it becomes clear that the current format is readily scalable for additional genes/strains.

After screening the strains available in our group, we found, in general, that 10 out of the 16 targets, or 63% were recovered in reasonable amounts. In at least four cases, the recovery was observed in multiple strains, while leastwise in three cases, a single strain was found to be more efficient in recovering a target in the soluble form (**T7, T13**, and **T14**). In the other cases, including the two full-length glycosyltransferases, none of the strains used here was found to recover the target in the soluble form.

To be able to test several different targets in a batch experiment, we missed some optimization procedures that could enhance our recovering rate. For example, the Lemo21 strain can fine-tune the expression level of toxic protein by the titratable regulation of the T7 lysozyme (T7L) expression using L-rhamnose [11,12]. In addition, since one would expect to observe some variation in protein expression among cell colonies/clones, it would be desirable to test more than one clone per target/strain. However, the number of experiments to run increases rapidly by testing all these variables. For this study, just a single clone was used, considering that a finer screening of colonies for cell expression could be done in a further stage.

pT-GroE, a strain that co-expresses a molecular chaperone to assist the proper protein folding, was found to result in a good recovering rate as well as a good contrast from the target protein to the basal protein expression, making it very easy to follow the target protein being expressed. Similar to pT-GroE, the Arctic Express strain combines low temperature-adapted chaperones Cpn10 and Cpn60, similar to *E. coli* GroEL and GroES, to result in chaperon assisted and low-temperature protein expression [13]. This strategy was successful in several targets tested here and was also the strain demonstrating a higher recovering rate.

The recovering rate in any structural biology pipeline will always be highly dependent on the targets chosen in the project. In this sense, we have no aim to claim that a given commercial strain is better or worse than each other. Rather than that, we show that it might be worth having a suitable and scalable assay to screen different host strains for the expression of soluble protein in difficult cases. We easily foresee a scenario where proteins that were not recovered in the soluble fraction after a ‘*standard’* protocol can be re-assayed for a ‘*salvage’* pipeline, where different strains are tested for the expression. The experimental cost of this effort includes the transformation of the existing vector in different cell strains and the implementation of a protocol for expression and purification that is scalable and may be implemented in HT format. Protein targets that cannot be recovered in this salvage pipeline are still suitable for changing the construction or even going for a eukaryotic cell factory as a host for heterologous expression. In these cases, the experimental costs are slightly higher and involve going back to the construct and primer design, gene cloning, and testing for expression again.

## 5. Conclusion

In conclusion, here we demonstrated that 10 out of 16 ‘difficult targets’ could be recovered in the soluble fraction after testing a set of different *E. coli* strains. This easy-to-implement assay is scalable for a different number of genes and/or strains. We found that the screening may represent an affordable salvage pipeline for the recovery of target proteins which complements a typical pipeline for protein expression and purification in structural biology efforts.

## Acknowledgments

The authors thank to the members of the Molecular Biotecnology Group @ IFSC/USP and, in particular, for our technical staff: Maria Auxiliadora Santos, João Possatto and Josimar Sartori. The authors also thank the financial support from the Fundação de Apoio À Pesquisa do Estado de São Paulo (FAPESP) through grants, 2015/26722-8, 2017/03966-4, 2017/24901-8, 2018/21213-6, 2019/20219-3 and 2020/03983-9 and from the Conselho Nacional de Desenvolvimento Científico e Tecnológico (CNPq) through grants 303165/2018-9, 485950/2013-8 and 476606/2010-1. This study was also financed in part by the Coordenação de Aperfeiçoamento de Pessoal de Nível Superior - Brasil (CAPES) - Finance Code 001.

## References

1. Beddell CR, Goodford PJ, Norrington FE, Wilkinson S, Wootton R. COMPOUNDS DESIGNED TO FIT A SITE OF KNOWN STRUCTURE IN HUMAN HAEMOGLOBIN. Br J Pharmacol. 1976;57: 201– 209. doi:10.1111/j.1476-5381.1976.tb07468.x

2. Rosano GL, Ceccarelli EA. Microbial Cell Factories Rare codon content affects the solubility of recombinant proteins in a codon bias-adjusted Escherichia coli strain. 2009; doi:10.1186/1475-2859-8-41

3. Kaur J, Kumar A, Kaur J. Strategies for optimization of heterologous protein expression in E. coli: Roadblocks and reinforcements. International Journal of Biological Macromolecules. Elsevier B.V.; 2018. pp. 803–822. doi:10.1016/j.ijbiomac.2017.08.080

4. Sonoda MT, Godoy AS, Pellegrini VOA, Kadowaki MAS, Nascimento AS, Polikarpov I. Structure and dynamics of Trichoderma harzianum Cel7B suggest molecular architecture adaptations required for a wide spectrum of activities on plant cell wall polysaccharides. Biochim Biophys Acta - Gen Subj. Elsevier; 2019;1863: 1015–1026. doi:10.1016/j.bbagen.2019.03.013

5. Camilo CM, Polikarpov I. High-throughput cloning, expression and purification of glycoside hydrolases using Ligation-Independent Cloning (LIC). Protein Expr Purif. Elsevier Inc.; 2014;99: 35–42. doi:10.1016/j.pep.2014.03.008

6. Eschenfeldt WH, Maltseva N, Stols L, Donnelly MI, Gu M, Nocek B, et al. Cleavable C-terminal His-tag vectors for structure determination. J Struct Funct Genomics. 2010;11: 31–39. doi:10.1007/s10969-010-9082-y

7. Stols L, Gu M, Dieckman L, Raffen R, Collart FR, Donnelly MI. A new vector for high-throughput, ligation-independent cloning encoding a tobacco etch virus protease cleavage site. Protein Expr Purif. 2002;25: 8–15. doi:10.1006/prep.2001.1603

8. Eschenfeldt WH, Lucy S, Millard CS, Joachimiak A, Mark ID. A family of LIC vectors for high-throughput cloning and purification of proteins. Methods Mol Biol. 2009;498: 105–15. doi:10.1007/978-1-59745-196-3_7

9. Studier FW. Protein production by auto-induction in high-density shaking cultures. 2005; doi:10.1016/j.pep.2005.01.016

10. Schneider CA, Rasband WS, Eliceiri KW. NIH Image to ImageJ: 25 years of image analysis. Nat Methods. Nature Publishing Group; 2012;9: 671–675. doi:10.1038/nmeth.2089

11. Schlegel S, Löfblom J, Lee C, Hjelm A, Klepsch M, Strous M, et al. Optimizing membrane protein overexpression in the Escherichia coli strain Lemo21(DE3). J Mol Biol. Academic Press; 2012;423: 648–659. doi:10.1016/j.jmb.2012.07.019

12. Schlegel S, Rujas E, Ytterberg AJ, Zubarev RA, Luirink J, de Gier JW. Optimizing heterologous protein production in the periplasm of E. coli by regulating gene expression levels. Microb Cell Fact. BioMed Central; 2013;12: 24. doi:10.1186/1475-2859-12-24

13. Ferrer M, Chernikova TN, Yakimov MM, Golyshin PN, Timmis KN. Chaperonins govern growth of Escherichia coli at low temperatures. Nat Biotechnol. Springer Science and Business Media LLC; 2003;21: 1267–1267. doi:10.1038/nbt1103-1266b

14. Gasteiger E, Hoogland C, Gattiker A, Duvaud S, Wilkins MR, Appel RD, et al. Protein Identification and Analysis Tools on the ExPASy Server. The Proteomics Protocols Handbook. Totowa, NJ: Humana Press; 2005. pp. 571–607. doi:10.1385/1-59259-890-0:571

