## Supplementary figures and images for "A Scalable Screening of *E. coli* Strains for Recombinant Protein Expression"

### Supplemental Figure 1a

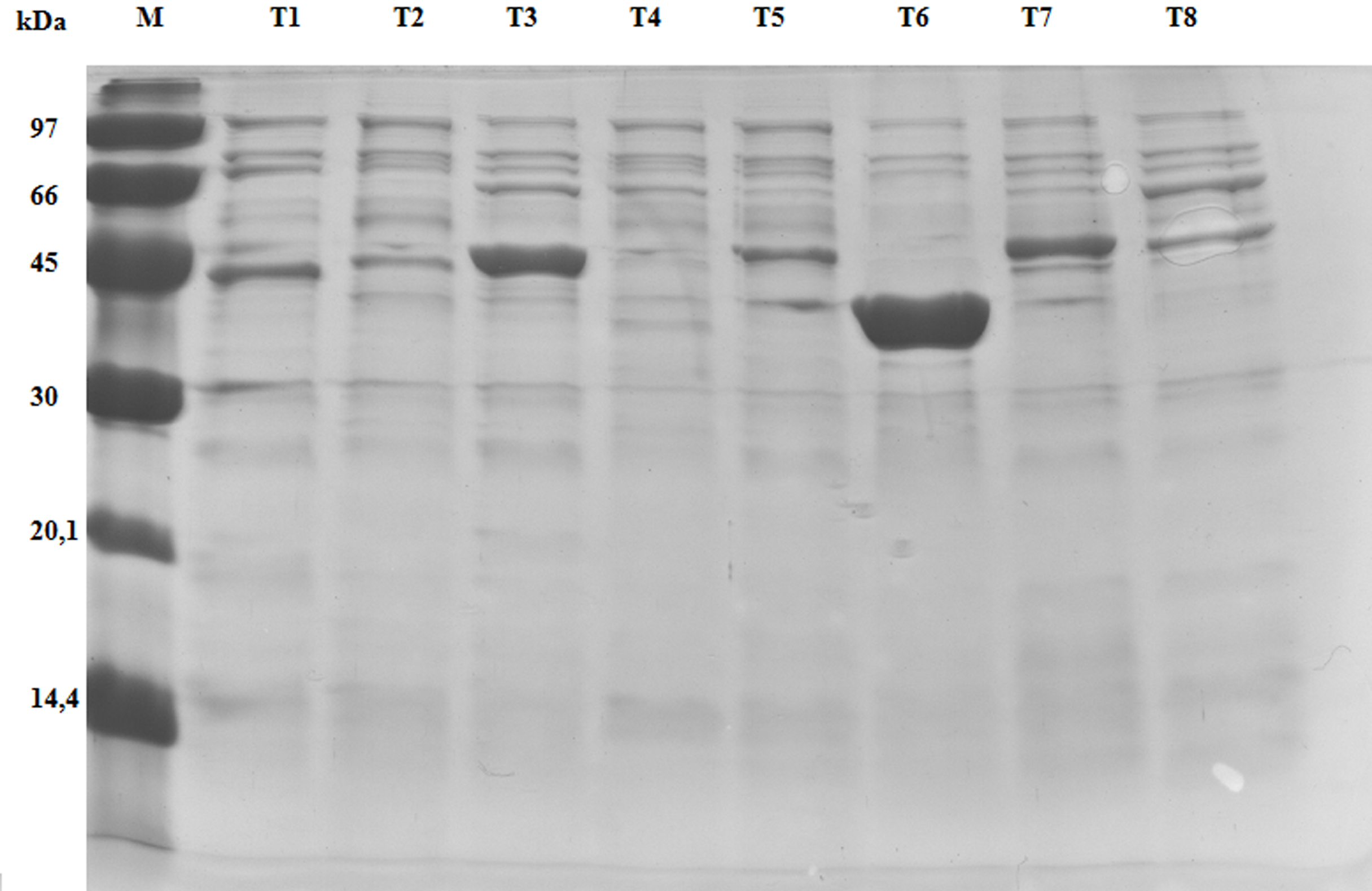

### Supplemental Figure 1b

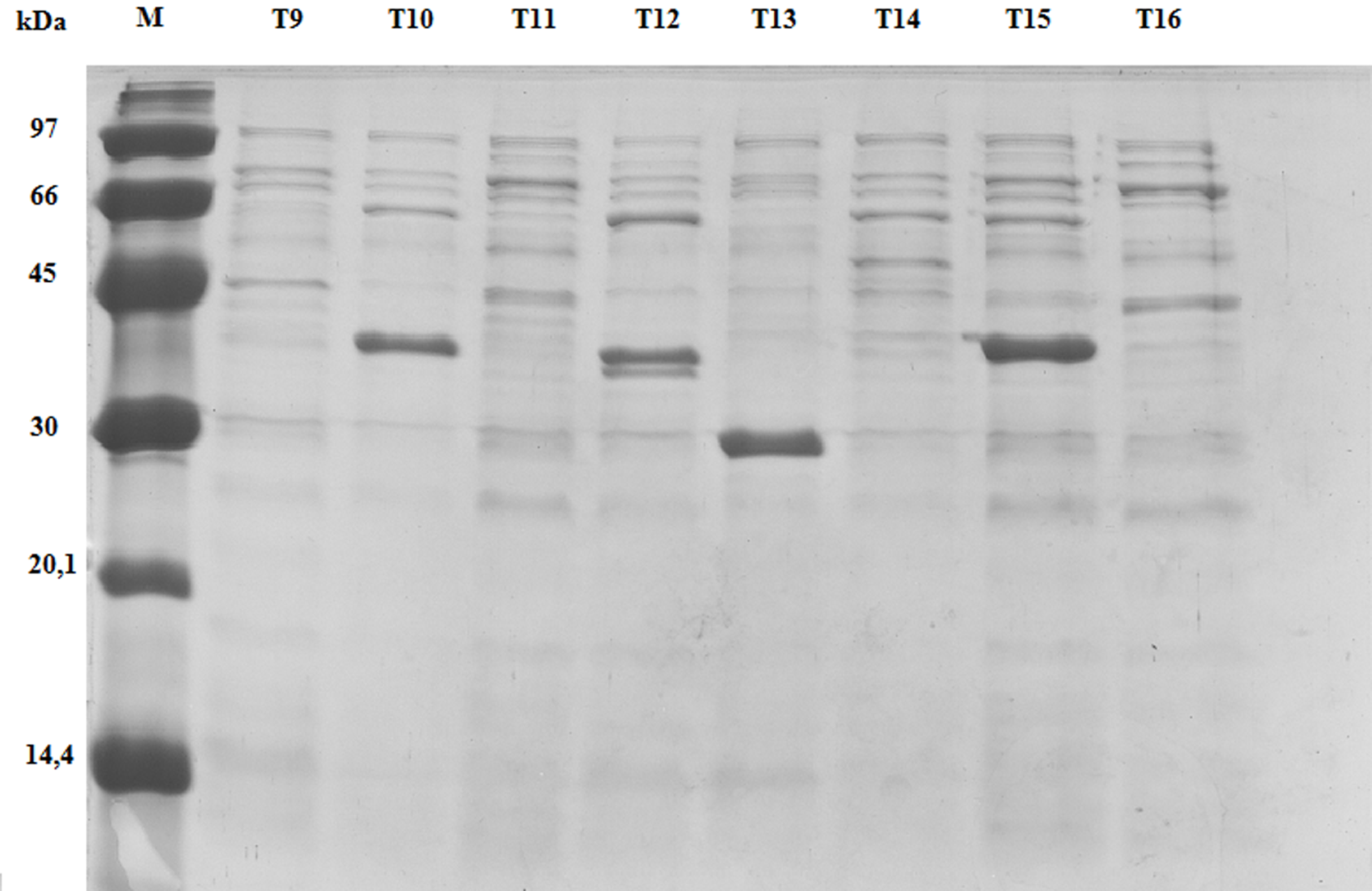

### Supplemental Figure 2a

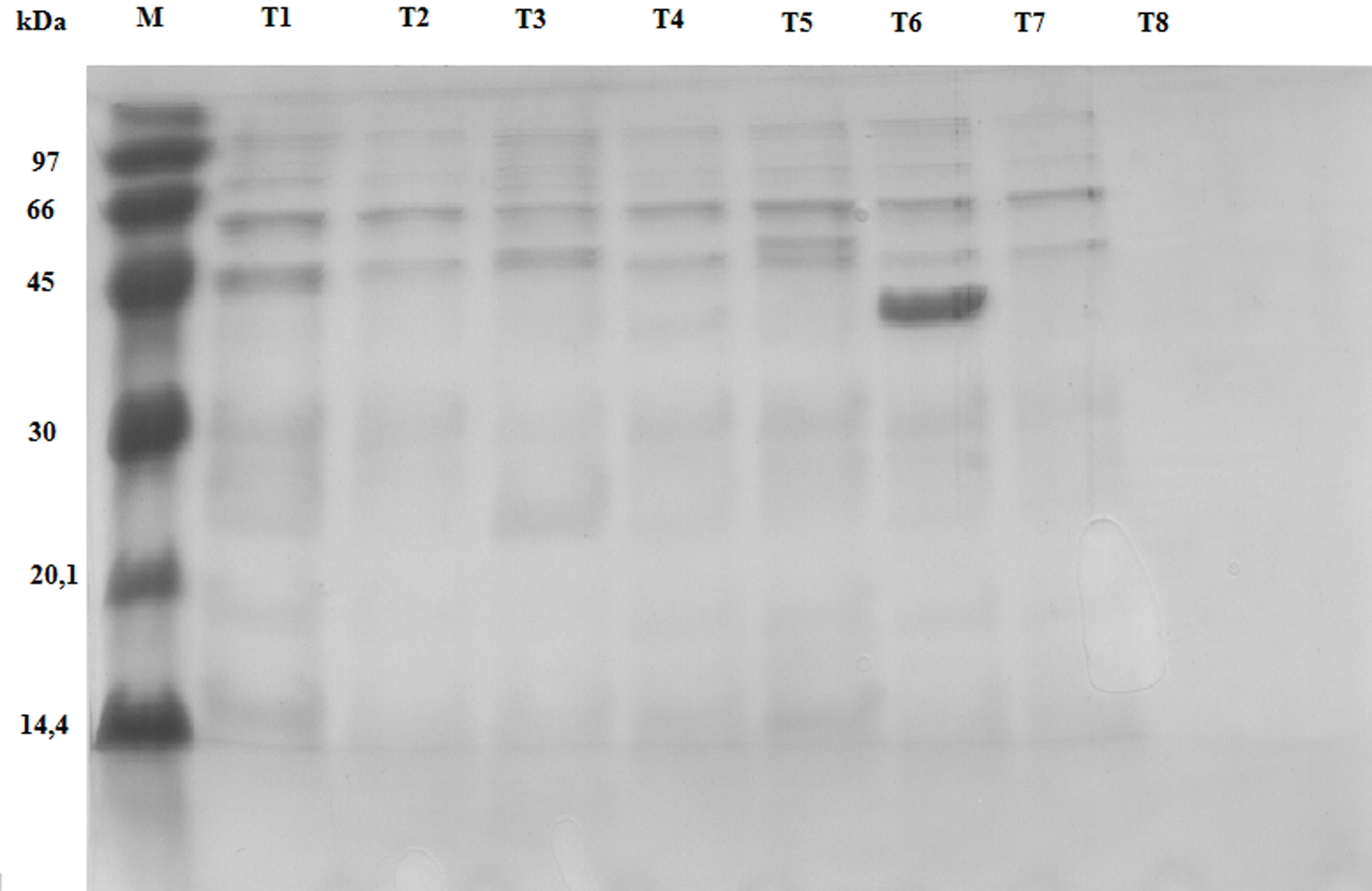

### Supplemental Figure 2b

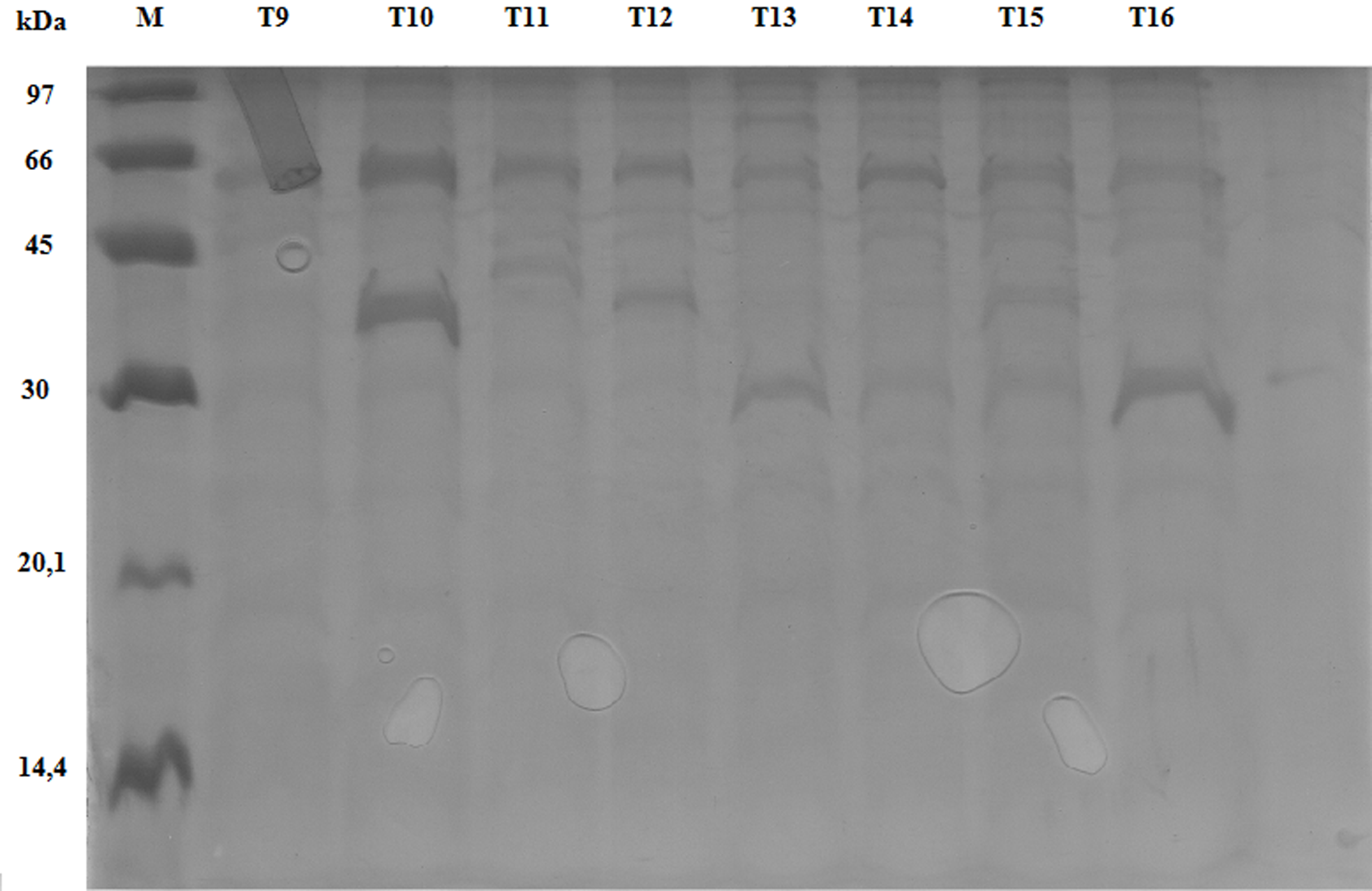

### Supplemental Figure 3a

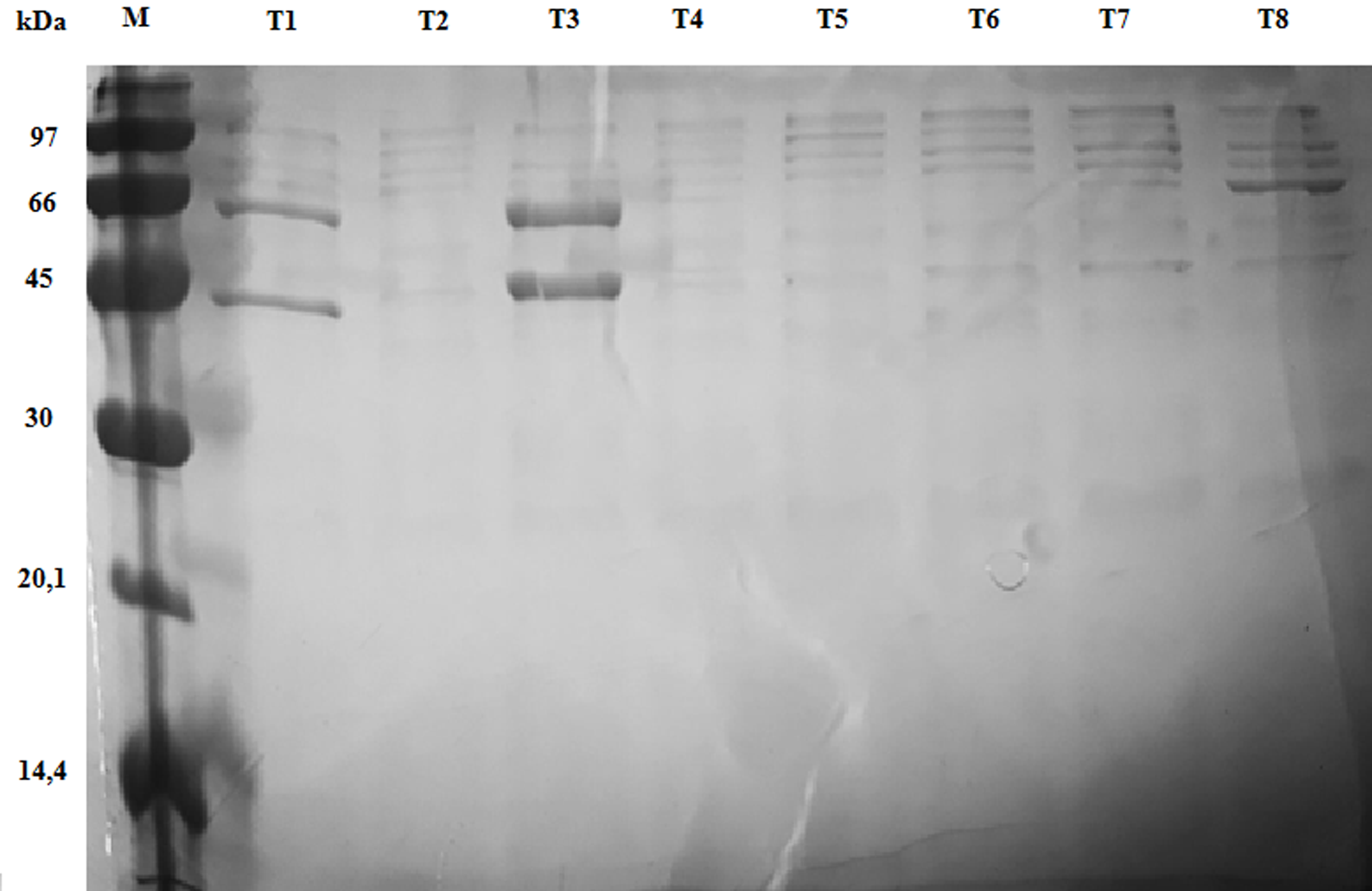

### Supplemental Figure 3b

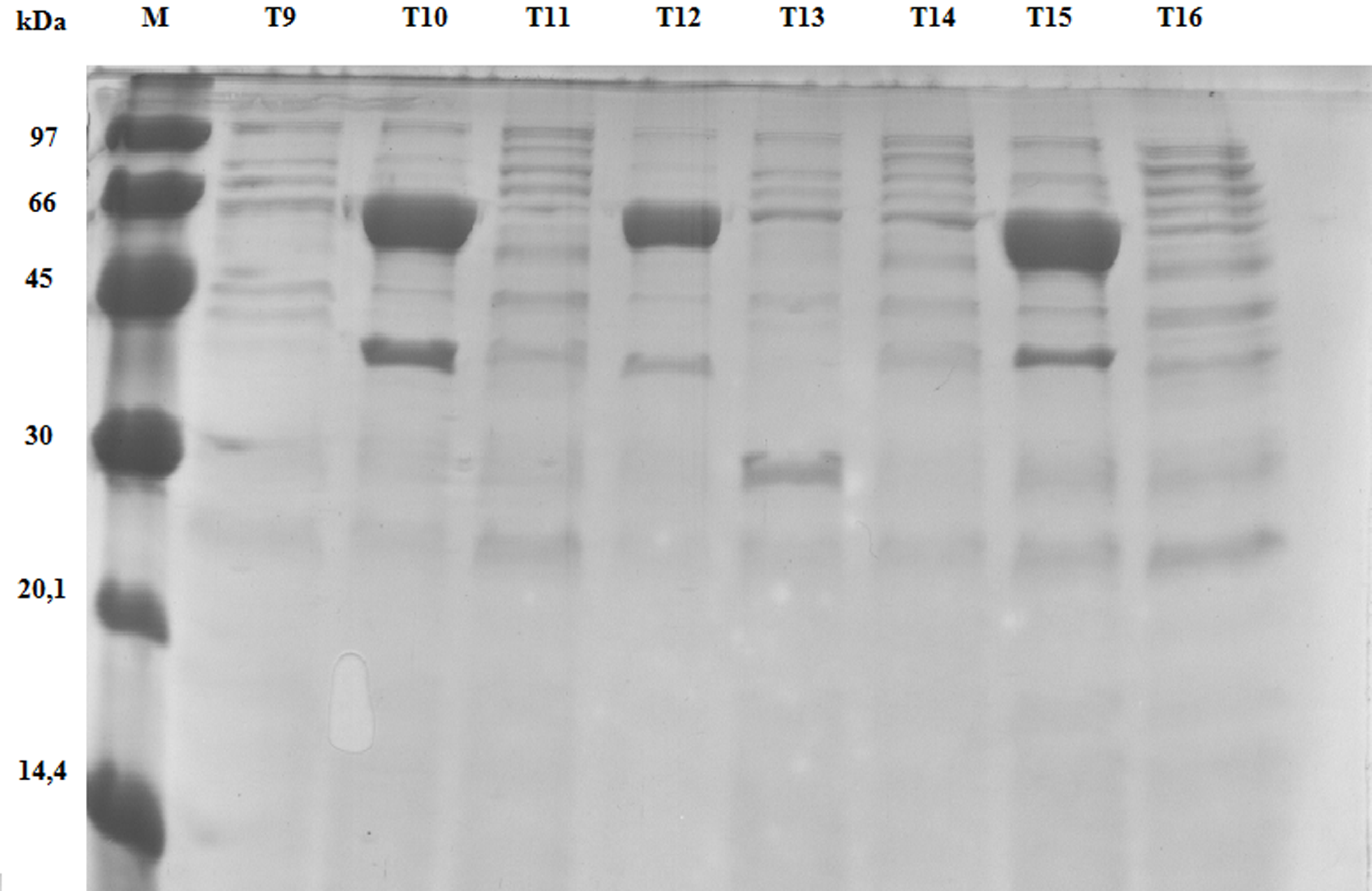

### Supplemental Figure 4a

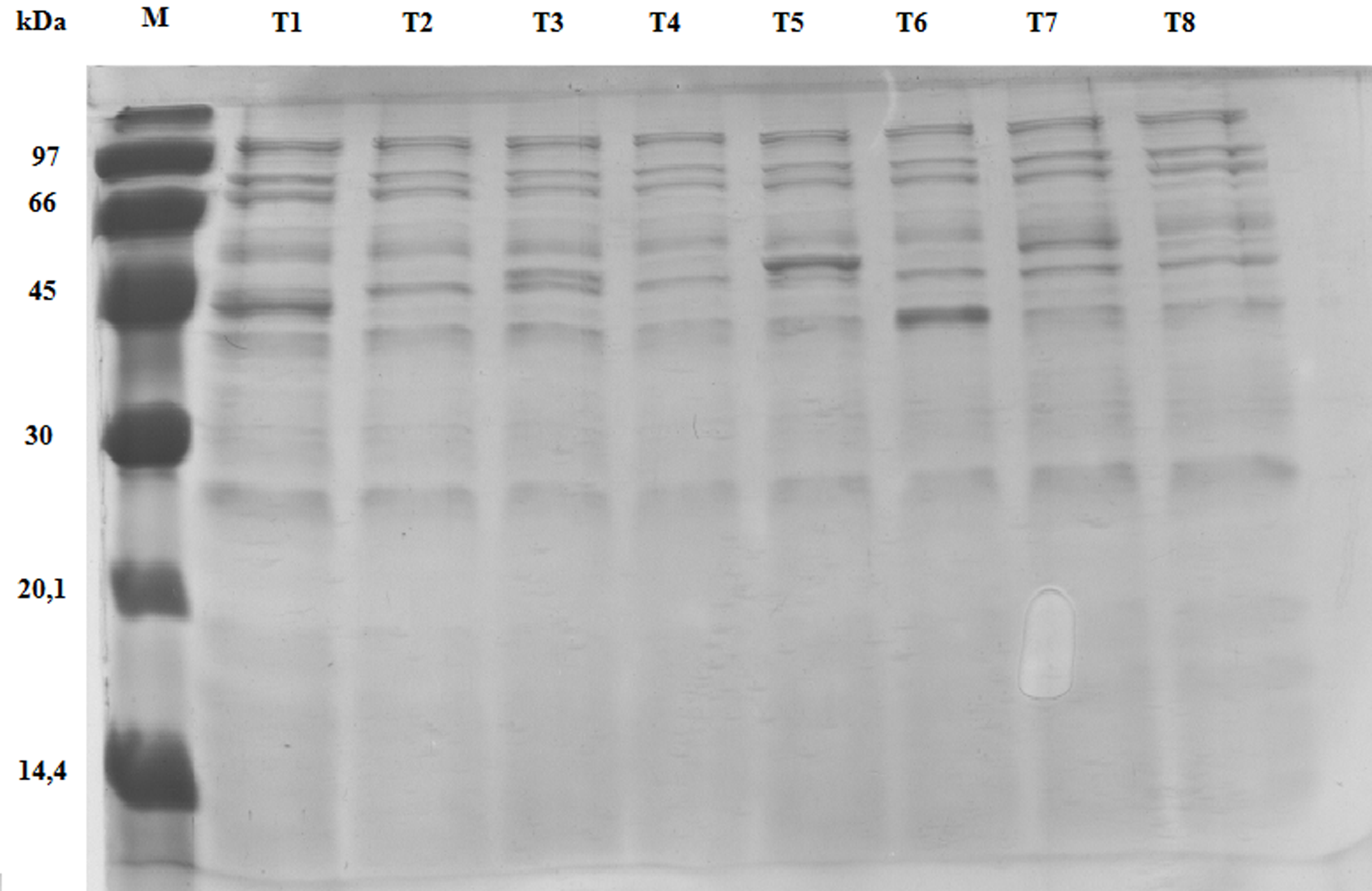

### Supplemental Figure 4b

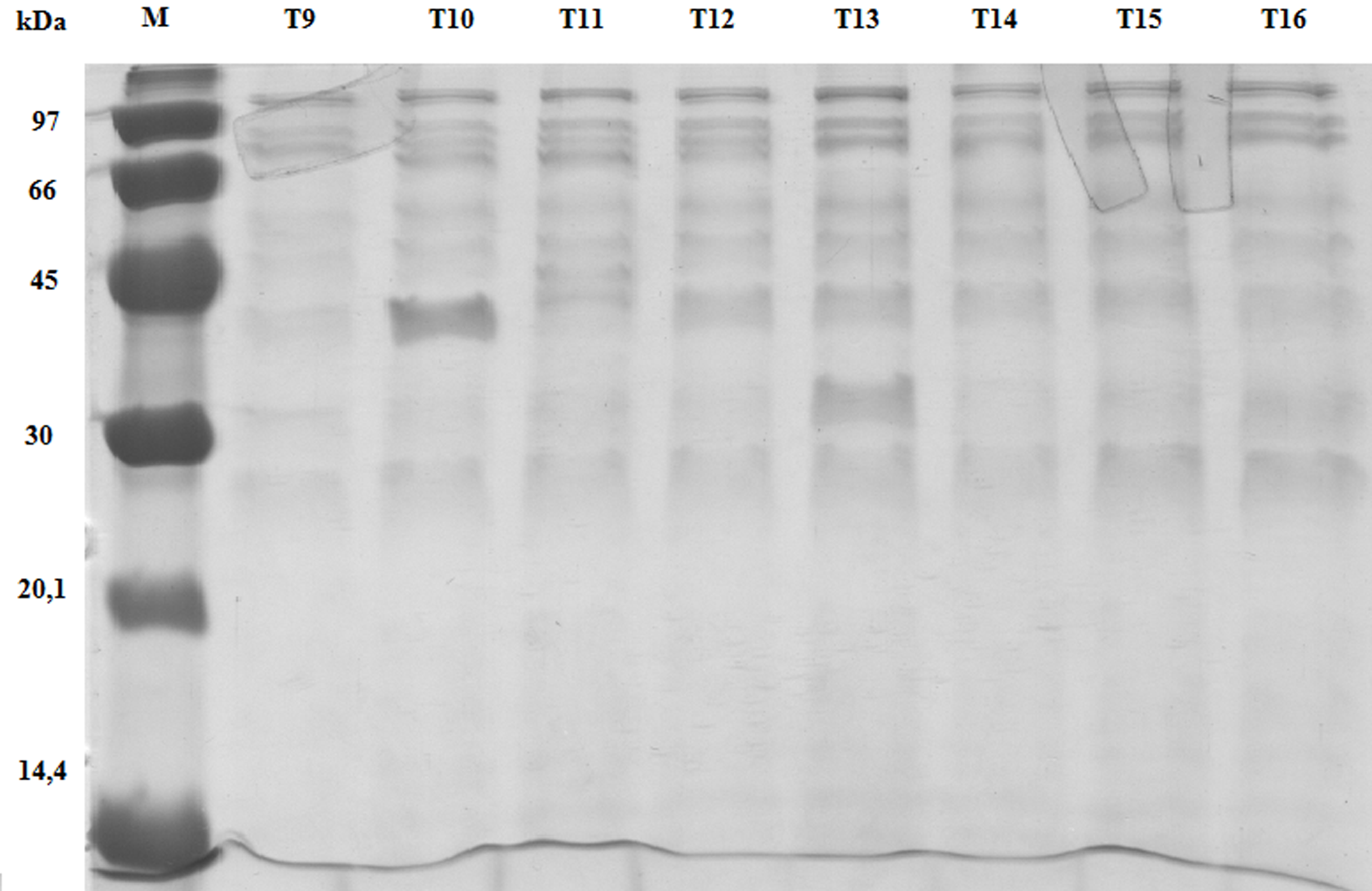
